# An ATAC-seq atlas of chromatin accessibility in mouse tissues

**DOI:** 10.1101/582171

**Authors:** Chuanyu Liu, Mingyue Wang, Xiaoyu Wei, Liang Wu, Jiangshan Xu, Xi Dai, Jun Xia, Mengnan Cheng, Yue Yuan, Pengfan Zhang, Jiguang Li, Taiqing Feng, Ao Chen, Wenwei Zhang, Fang Chen, Zhouchun Shang, Xiuqing Zhang, Brock A. Peters, Longqi Liu

## Abstract

The Assay for Transposase-Accessible Chromatin using sequencing (ATAC-seq) is a fundamental epigenomics approach and has been widely used in profiling the chromatin accessibility dynamics in multiple species. A comprehensive reference of ATAC-seq datasets for mammalian tissues is important for the understanding of regulatory specificity and developmental abnormality caused by genetic or environmental alterations. Here, we report an adult mouse ATAC-seq atlas by producing a total of 66 ATAC-seq profiles from 20 primary tissues of both male and female mice. The ATAC-seq read enrichment, fragment size distribution, and reproducibility between replicates demonstrated the high quality of the full dataset. We identified a total of 296,574 accessible elements, of which 26,916 showed tissue-specific accessibility. Further, we identified key transcription factors specific to distinct tissues and found that the enrichment of each motif reflects the developmental similarities across tissues. In summary, our study provides an important resource on the mouse epigenome and will be of great importance to various scientific disciplines such as development, cell reprogramming, and genetic disease.

## Background & Summary

Although most of the protein-coding genes in human and model animals such as mouse have been extensively annotated, vast regions of the genome are noncoding sequences (e.g., roughly 98% of the human genome) and still poorly understood^1,2^. During the last decade, the development of next-generation sequencing (NGS) based epigenomics techniques (e.g., ChIP-seq and DNase-seq) have significantly facilitated the identification of functional genomic regions^3^. For example, by comparing the histone modifications and transcription factor (TF) binding patterns throughout the mouse genome in a wide spectrum of tissues and cell types, Yue *et al*.^4,5^ have made significant progress towards a comprehensive catalog of potential functional elements in the mouse genome. So far, the international human epigenome consortium (IHEC), including ENCODE and the NIH Roadmap epigenomics projects, have profiled thousands of epigenomes including DNA methylation, genome-wide binding of TFs, histone modifications, and chromatin accessibility. This has resulted in the discovery of over 5 million *cis*-regulatory elements (CREs) in the human genome^6–8^. These data resources have created an important baseline for further study of diverse biological processes, such as development, cell reprogramming, and human disease^9–13^.

The accessibility of CREs, which is important for switching on and off gene expression^14^, is strongly associated with transcriptional activity. To date, detection of DNase I hypersensitive sites (DHSs) within chromatin by DNase-seq has been extensively used to map accessible genomic regions in diverse organisms including the laboratory mouse^5^. In 2013, Buenrostro *et al*.^15^ reported an alternative approach, termed ATAC-seq, for fast and sensitive profiling of chromatin accessibility by direct transposition of native chromatin within the nucleus. This method, in comparison to DNase-seq, requires a significantly lower input of cells (only 500-50,000) and a shorter period to process samples^16^. Moreover, ATAC-seq has been applied to single cells through various methods^17–20^, enabling the investigation of regulatory heterogeneity within complex tissues. As such, ATAC-seq has demonstrated great potential to be a leading method in assaying accessible chromatin genome-wide.

The sequence preference of both DNase I and Tn5 enzymes produced distinct but inevitable biases in DNase-seq and ATAC-seq^21^, making it impractical to directly compare datasets generated from the two methods. Therefore, although the DNase-seq atlas of adult mouse tissues has been published^5^, a baseline of chromatin accessible regions generated from ATAC-seq is still important for ATAC-seq based studies. Here, we applied Omni-ATAC-seq^22^, an approach that enables profiling of accessible chromatin from frozen tissues, to the generation of 66 chromatin accessibility datasets from 20 different tissues derived from both adult male and female C57BL/6J mice (Fig. 1a). Systematic analysis of the dataset identified a total of 296,574 accessible elements, of which 26,916 showed highly tissue-specific patterns. We further predicted TFs specific to distinct tissues and importantly, many of these have been validated by previous studies^23–27^. In this study, we provide a valuable resource, which can be used to elucidate transcriptional regulation and may further help understand diseases caused by regulatory dysfunction.

**Figure 1.**
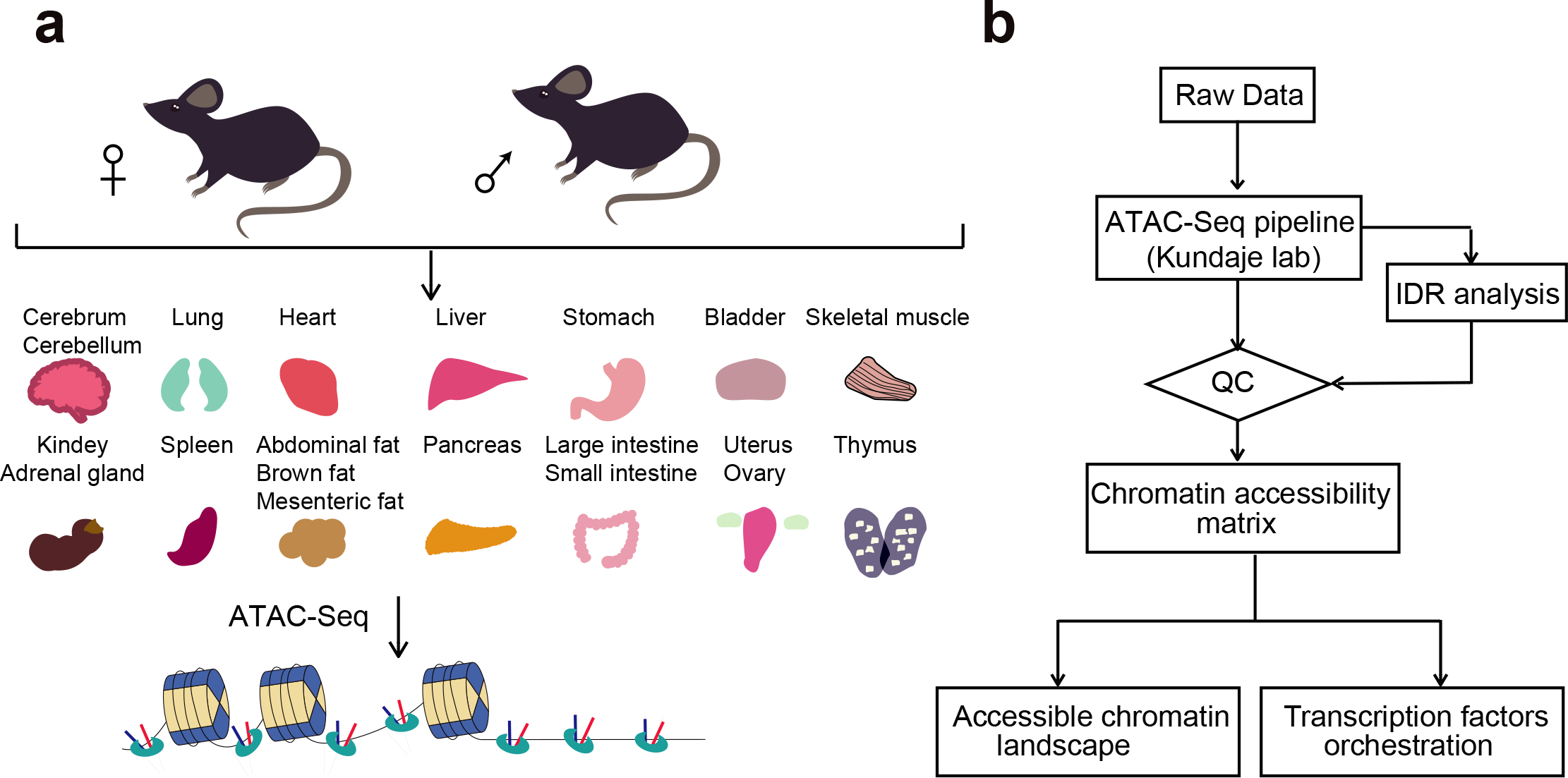
Overview of the experimental and data analysis workflow. (**a**) 20 different tissues from adult mice were collected for ATAC-seq profiling. (**b**) The analysis workflow for ATAC-seq profiles.

## Methods

### Sample collection

All relevant procedures involving animals were approved by the Institutional Review Board on Ethics Committee of BGI (Permit No. BGI-R085-1). C57BL/6J male and female mice were purchased from Beijing Vital River Laboratory Animal Technology Co., Ltd (Beijing, China). 8-week old mice were used for this study. Mice were housed under standard conditions of a specific pathogen-free, temperature-controlled environment with a 12-h day/night cycle^28^. The mice were sacrificed by cervical dislocation. Whole organs were extracted and cut into 2-3 pieces, respectively (50-200 mg/piece). Each sample was then quickly frozen in liquid nitrogen and stored at −80 °C until nuclei extraction was performed. In this study, we used 20 different organs or tissues, including adrenal gland, bladder, brain (cerebrum and cerebellum), fat (abdominal, brown and mesenteric), heart, intestine (large and small), kidney, liver, lung, ovary, pancreas, skeletal muscle, spleen, stomach, thymus, and uterus (as listed in Table 1).

**Table 1.** Tissue and corresponding mouse and sample IDs.

| Sample ID | Strain | Serial | Gender | Tissue | Replicate |
| --- | --- | --- | --- | --- | --- |
| ATAC-1 | C57BL/6J | 1 | Female | Adrenal gland | 1 |
| ATAC-2 | C57BL/6J | 1 | Female | Adrenal gland | 2 |
| ATAC-3 | C57BL/6J | 1 | Female | Cerebrum | 1 |
| ATAC-4 | C57BL/6J | 1 | Female | Cerebrum | 2 |
| ATAC-5 | C57BL/6J | 1 | Female | Cerebellum | 1 |
| ATAC-5 | C57BL/6J | 1 | Female | Cerebellum | 2 |
| ATAC-7 | C57BL/6J | 1 | Female | Abdominal fat | 1 |
| ATAC-8 | C57BL/6J | 1 | Female | Abdominal fat | 2 |
| ATAC-9 | C57BL/6J | 1 | Female | Brown fat | 1 |
| ATAC-10 | C57BL/6J | 1 | Female | Brown fat | 2 |
| ATAC-11 | C57BL/6J | 1 | Female | Mesenteric fat | 1 |
| ATAC-12 | C57BL/6J | 1 | Female | Mesenteric fat | 2 |
| ATAC-13 | C57BL/6J | 1 | Female | Heart | 1 |
| ATAC-14 | C57BL/6J | 1 | Female | Heart | 2 |
| ATAC-15 | C57BL/6J | 1 | Female | Kidney | 1 |
| ATAC-16 | C57BL/6J | 1 | Female | Kidney | 2 |
| ATAC-17 | C57BL/6J | 1 | Female | Liver | 1 |
| ATAC-18 | C57BL/6J | 1 | Female | Liver | 2 |
| ATAC-19 | C57BL/6J | 1 | Female | Lung | 1 |
| ATAC-20 | C57BL/6J | 1 | Female | Lung | 2 |
| ATAC-21 | C57BL/6J | 1 | Female | Ovary | 1 |
| ATAC-22 | C57BL/6J | 1 | Female | Ovary | 2 |
| ATAC-23 | C57BL/6J | 1 | Female | Pancreas | 1 |
| ATAC-24 | C57BL/6J | 1 | Female | Pancreas | 2 |
| ATAC-25 | C57BL/6J | 1 | Female | Skeletal muscle | 1 |
| ATAC-26 | C57BL/6J | 1 | Female | Skeletal muscle | 2 |
| ATAC-27 | C57BL/6J | 1 | Female | Spleen | 1 |
| ATAC-28 | C57BL/6J | 1 | Female | Spleen | 2 |
| ATAC-29 | C57BL/6J | 1 | Female | Thymus | 1 |
| ATAC-30 | C57BL/6J | 1 | Female | Thymus | 2 |
| ATAC-31 | C57BL/6J | 1 | Female | Uterus | 1 |
| ATAC-32 | C57BL/6J | 1 | Female | Uterus | 2 |
| ATAC-33 | C57BL/6J | 2 | Male | Adrenal gland | 1 |
| ATAC-34 | C57BL/6J | 2 | Male | Adrenal gland | 2 |
| ATAC-35 | C57BL/6J | 2 | Male | Bladder | 1 |
| ATAC-36 | C57BL/6J | 2 | Male | Bladder | 2 |
| ATAC-37 | C57BL/6J | 2 | Male | Cerebrum | 1 |
| ATAC-38 | C57BL/6J | 2 | Male | Cerebrum | 2 |
| ATAC-39 | C57BL/6J | 2 | Male | Cerebellum | 1 |
| ATAC-40 | C57BL/6J | 2 | Male | Cerebellum | 2 |
| ATAC-41 | C57BL/6J | 2 | Male | Brown fat | 1 |
| ATAC-42 | C57BL/6J | 2 | Male | Brown fat | 2 |
| ATAC-43 | C57BL/6J | 2 | Male | Mesenteric fat | 1 |
| ATAC-44 | C57BL/6J | 2 | Male | Mesenteric fat | 2 |
| ATAC-45 | C57BL/6J | 2 | Male | Heart | 1 |
| ATAC-46 | C57BL/6J | 2 | Male | Heart | 2 |
| ATAC-47 | C57BL/6J | 2 | Male | Large intestine | 1 |
| ATAC-48 | C57BL/6J | 2 | Male | Large intestine | 2 |
| ATAC-49 | C57BL/6J | 2 | Male | Small intestine | 1 |
| ATAC-50 | C57BL/6J | 2 | Male | Small intestine | 2 |
| ATAC-51 | C57BL/6J | 2 | Male | Kidney | 1 |
| ATAC-52 | C57BL/6J | 2 | Male | Kidney | 2 |
| ATAC-53 | C57BL/6J | 2 | Male | Liver | 1 |
| ATAC-54 | C57BL/6J | 2 | Male | Liver | 2 |
| ATAC-55 | C57BL/6J | 2 | Male | Lung | 1 |
| ATAC-56 | C57BL/6J | 2 | Male | Lung | 2 |
| ATAC-57 | C57BL/6J | 2 | Male | Pancreas | 1 |
| ATAC-58 | C57BL/6J | 2 | Male | Pancreas | 2 |
| ATAC-59 | C57BL/6J | 2 | Male | Skeletal muscle | 1 |
| ATAC-60 | C57BL/6J | 2 | Male | Skeletal muscle | 2 |
| ATAC-61 | C57BL/6J | 2 | Male | Spleen | 1 |
| ATAC-62 | C57BL/6J | 2 | Male | Spleen | 2 |
| ATAC-63 | C57BL/6J | 2 | Male | Stomach | 1 |
| ATAC-64 | C57BL/6J | 2 | Male | Stomach | 2 |
| ATAC-65 | C57BL/6J | 2 | Male | Thymus | 1 |
| ATAC-66 | C57BL/6J | 2 | Male | Thymus | 2 |

### Library construction and sequencing

Tissues were homogenized in a 2 ml Dounce homogenizer (with a loose and then tight pestle) with 10-20 strokes in 2 ml of 1 X homogenization buffer on ice. 400 μl of this nuclei suspension was transferred to a round-bottom 2 ml Lo-Bind Eppendorf tube for density gradient centrifugation following the protocol by Corces *et al*.^22^. After centrifugation, the nuclei band (about 200 μl) was collected, stained with DAPI, and nuclei were counted. Approximately 20,000-100,000 nuclei were transferred into a fresh tube and diluted in 1 ml ATAC-RSB + 0.1% Tween-20 (Sigma-Aldrich, Darmstadt, Germany). Nuclei were centrifuged and the supernatant was carefully aspirated. Nuclei were treated in 50 μl transposition reaction mixture containing 10 mM TAPS-NaOH (pH 8.5), 5 mM MgCl_2_, 10% DMF, 2.5 μl of in-house Tn5 transposase (0.8 U/μl), 0.01% digitonin (Sigma-Aldrich, Darmstadt, Germany), 0.1% Tween-20, 31.5 μl of PBS, and 5 μl of nuclease-free water for 30 mins at 37 °C. Afterward, the DNA was purified with MinElute Purification Kit (Qiagen, Venlo, Netherlands) and amplified with primers containing barcodes, as previously described^22,29^.

All libraries were adapted for sequencing on the BGISEQ-500 platform^30^. In brief, the DNA concentration was determined by Qubit 3.0 (ThermoFisher, Waltham, MA). Pooled samples were used to make single-strand DNA circles (ssDNA circles). DNA nanoballs (DNBs) were generated from the ssDNA circles by rolling circle replication as previously described^30^. The DNBs were loaded onto patterned nano-arrays and sequenced on the BGISEQ-500 sequencing platform with paired end 50 base reads.

### Preprocessing of the ATAC-seq datasets

The ATAC-seq data were processed (trim, alignment, filter, and QC) using the ATAC-seq pipeline from the Kundaje lab^31,32^ (Table 2). The model-based analysis of ChIP-seq (MACS2)^33^ version 2.1.2 was used to identify the peak regions with options −B, −q 0.01 -- nomodel, −f BAM, −g mm. The Irreproducible Discovery Rate (IDR) method^34^ was used to identify reproducible peaks between two technical replicates (Fig. 1b). Only peaks reproducible between the two technical replicates were retained for downstream analyses. We established a standard peak set by merging all overlapping peaks. The number of raw reads mapped to each standard peak were counted using the *intersect* function of BedTools^35^ version 2.26.0. The raw count matrix^32^ was normalized by Reads Per Million mapped reads (RPM). Pearson correlation coefficients between technical or biological replicates across tissues were calculated based on the Log10 RPM matrix.

**Table 2.** ATAC-seq metadata and mapping statistics. *Mapped reads: total number of read minus number of unaligned read; *Usable reads: number of mapped read minus number of low mapping quality, duplicate and mitochondrial reads.

| Sample ID | Total Reads | Mapped Reads* | chrM Reads | Usable Reads* | Percentage of Usable Reads | TSS Enrichment | IDR Peaks |
| --- | --- | --- | --- | --- | --- | --- | --- |
| ATAC-1 | 98,303,382 | 96,954,685 | 23,288,521 | 16,098,792 | 16.38 | 17.21 | 23,296 |
| ATAC-2 | 148,416,230 | 146,416,001 | 25,259,955 | 24,542,678 | 16.54 | 17.83 | 23,296 |
| ATAC-3 | 150,254,776 | 148,205,698 | 1,773,531 | 116,676,430 | 77.65 | 18.35 | 93,546 |
| ATAC-4 | 132,170,414 | 130,467,930 | 1,749,710 | 108,546,212 | 82.13 | 19.17 | 93,546 |
| ATAC-5 | 44,251,998 | 43,732,156 | 254,745 | 35,315,756 | 79.81 | 13.90 | 32,337 |
| ATAC-5 | 180,555,698 | 178,313,881 | 2,085,323 | 136,439,604 | 75.57 | 14.51 | 32,337 |
| ATAC-7 | 170,643,536 | 166,598,249 | 1,204,915 | 61,090,112 | 35.80 | 13.03 | 28,827 |
| ATAC-8 | 165,896,236 | 135,431,464 | 1,175,420 | 61,493,964 | 37.07 | 12.63 | 28,827 |
| ATAC-9 | 134,023,704 | 129,656,384 | 14,098,291 | 49,198,612 | 36.71 | 16.75 | 66,620 |
| ATAC-10 | 160,942,400 | 156,016,214 | 13,541,729 | 59,910,266 | 37.22 | 16.71 | 66,620 |
| ATAC-11 | 157,502,400 | 154,297,472 | 1,228,888 | 93,224,318 | 59.19 | 8.10 | 34,851 |
| ATAC-12 | 162,615,494 | 159,625,229 | 1,252,759 | 103,453,218 | 63.62 | 7.44 | 34,851 |
| ATAC-13 | 238,728,090 | 229,371,653 | 14,280,715 | 116,609,704 | 48.85 | 7.82 | 38,875 |
| ATAC-14 | 268,182,264 | 259,270,286 | 18,497,755 | 132,829,548 | 49.53 | 7.11 | 38,875 |
| ATAC-15 | 90,269,090 | 88,799,485 | 1,190,916 | 67,126,022 | 74.36 | 13.93 | 57,348 |
| ATAC-16 | 108,421,198 | 106,752,478 | 1,697,128 | 77,533,954 | 71.51 | 13.54 | 57,348 |
| ATAC-17 | 137,779,770 | 135,420,287 | 647,765 | 103,420,718 | 75.06 | 7.56 | 49,444 |
| ATAC-18 | 138,236,344 | 135,957,141 | 603,586 | 103,420,718 | 74.81 | 7.24 | 49,444 |
| ATAC-19 | 181,243,756 | 178,738,131 | 1,480,812 | 135,096,206 | 74.54 | 12.98 | 74,408 |
| ATAC-20 | 123,570,800 | 121,933,521 | 1,153,158 | 96,071,108 | 77.75 | 13.55 | 74,408 |
| ATAC-21 | 110,371,234 | 109,170,446 | 3,960,686 | 63,119,950 | 57.19 | 13.61 | 22,300 |
| ATAC-22 | 39,326,846 | 38,875,707 | 1,497,720 | 24,585,734 | 62.52 | 17.13 | 22,300 |
| ATAC-23 | 35,838,048 | 34,832,716 | 153,844 | 25,314,950 | 70.64 | 12.88 | 37,012 |
| ATAC-24 | 107,196,410 | 104,421,905 | 523,370 | 76,018,760 | 70.92 | 12.55 | 37,012 |
| ATAC-25 | 133,254,164 | 131,716,403 | 2,973,082 | 63,522,884 | 47.67 | 6.39 | 16,386 |
| ATAC-26 | 166,638,990 | 164,664,788 | 3,405,553 | 75,588,356 | 45.36 | 6.04 | 16,386 |
| ATAC-27 | 94,226,804 | 93,372,274 | 697,988 | 58,659,642 | 62.25 | 7.55 | 18,659 |
| ATAC-28 | 106,958,878 | 106,110,973 | 727,572 | 74,085,722 | 69.27 | 7.75 | 18,659 |
| ATAC-29 | 70,619,572 | 69,042,929 | 629,068 | 49,965,476 | 70.75 | 11.87 | 35,092 |
| ATAC-30 | 89,598,654 | 87,444,974 | 844,626 | 62,271,590 | 69.50 | 11.50 | 35,092 |
| ATAC-31 | 66,931,470 | 66,424,809 | 200,619 | 50,479,508 | 75.42 | 5.71 | 12,249 |
| ATAC-32 | 90,829,474 | 74,837,342 | 286,587 | 68,853,698 | 75.81 | 6.24 | 12,249 |
| ATAC-33 | 98,233,400 | 94,112,150 | 4,507,302 | 23,763,636 | 24.19 | 15.52 | 24,953 |
| ATAC-34 | 214,082,112 | 198,861,218 | 14,028,173 | 40,427,112 | 18.88 | 21.83 | 24,953 |
| ATAC-35 | 84,715,186 | 83,692,447 | 5,340,505 | 39,154,858 | 46.22 | 9.15 | 21,940 |
| ATAC-36 | 212,395,310 | 209,869,802 | 16,551,961 | 79,600,948 | 37.48 | 9.98 | 21,940 |
| ATAC-37 | 235,561,686 | 191,218,996 | 4,553,350 | 155,453,764 | 65.99 | 25.43 | 117,909 |
| ATAC-38 | 191,218,996 | 188,304,479 | 3,834,757 | 130,871,008 | 68.44 | 25.58 | 117,909 |
| ATAC-39 | 53,952,860 | 53,152,913 | 981,941 | 41,135,886 | 76.24 | 15.42 | 41,280 |
| ATAC-40 | 158,893,266 | 156,816,876 | 2,293,222 | 107,582,560 | 67.71 | 19.42 | 41,280 |
| ATAC-41 | 173,258,824 | 166,021,269 | 19,132,852 | 24,004,880 | 13.85 | 17.70 | 43,306 |
| ATAC-42 | 136,328,522 | 132,274,503 | 11,522,990 | 45,461,724 | 33.35 | 17.81 | 43,306 |
| ATAC-43 | 155,269,780 | 151,746,361 | 1,090,769 | 100,616,020 | 64.80 | 8.65 | 41,494 |
| ATAC-44 | 202,037,768 | 198,044,341 | 1,146,229 | 136,190,834 | 67.41 | 9.22 | 41,494 |
| ATAC-45 | 157,868,844 | 154,946,940 | 21,384,649 | 61,621,604 | 39.03 | 8.34 | 31,248 |
| ATAC-46 | 136,265,518 | 133,201,038 | 13,707,672 | 55,085,754 | 40.43 | 8.77 | 31,248 |
| ATAC-47 | 125,102,346 | 123,612,014 | 6,647,001 | 78,928,576 | 63.09 | 9.93 | 54,282 |
| ATAC-48 | 59,245,938 | 58,534,297 | 3,079,504 | 39,391,956 | 66.49 | 9.89 | 54,282 |
| ATAC-49 | 119,605,280 | 118,172,060 | 549,819 | 83,043,494 | 69.43 | 12.31 | 30,671 |
| ATAC-50 | 126,897,090 | 125,503,611 | 556,836 | 90,393,490 | 71.23 | 11.70 | 30,671 |
| ATAC-51 | 75,598,616 | 74,377,104 | 2,570,404 | 54,471,070 | 72.05 | 19.32 | 74,760 |
| ATAC-52 | 128,285,640 | 126,411,530 | 4,825,675 | 87,433,334 | 68.16 | 17.13 | 74,760 |
| ATAC-53 | 206,730,874 | 202,279,412 | 2,800,887 | 146,461,186 | 70.85 | 13.31 | 78,775 |
| ATAC-54 | 194,636,430 | 190,999,768 | 3,353,166 | 140,032,146 | 71.95 | 13.07 | 78,775 |
| ATAC-55 | 108,547,670 | 107,297,270 | 1,127,842 | 78,808,372 | 72.60 | 13.78 | 64,002 |
| ATAC-56 | 156,741,782 | 155,063,216 | 1,616,305 | 113,767,728 | 72.58 | 13.15 | 64,002 |
| ATAC-57 | 91,983,866 | 89,773,673 | 819,185 | 65,257,644 | 70.94 | 17.81 | 54,658 |
| ATAC-58 | 238,980,976 | 232,982,918 | 3,255,911 | 162,801,402 | 68.12 | 16.01 | 54,658 |
| ATAC-59 | 117,557,552 | 115,960,450 | 1,920,255 | 44,835,398 | 38.14 | 6.32 | 14,722 |
| ATAC-60 | 214,824,060 | 211,626,739 | 2,469,962 | 42,681,444 | 19.87 | 9.52 | 14,722 |
| ATAC-61 | 96,385,558 | 95,252,958 | 708,544 | 63,986,222 | 66.39 | 11.09 | 21,189 |
| ATAC-62 | 99,268,666 | 98,104,994 | 736,929 | 67,856,328 | 68.36 | 11.95 | 21,189 |
| ATAC-63 | 146,138,130 | 144,325,578 | 1,901,264 | 96,815,264 | 66.25 | 10.05 | 34,443 |
| ATAC-64 | 53,691,468 | 52,961,106 | 1,045,366 | 37,522,232 | 69.88 | 13.43 | 34,443 |
| ATAC-65 | 108,290,790 | 106,397,126 | 1,095,779 | 78,782,088 | 72.75 | 14.54 | 42,037 |
| ATAC-66 | 131,298,716 | 128,823,646 | 1,286,797 | 93,295,128 | 71.06 | 14.16 | 42,037 |

### Identification of tissue-specific chromatin accessible regions

We used a strategy described previously based on the Shannon entropy to compute a tissue specificity index for each peak^4,36,37^. Specifically, for each peak, we defined its relative accessibility in a tissue type i as Ri = Ei / ΣE, where Ei is the RPM value for the peak in the tissue i ΣE is the sum of RPM values in all tissues, and N is the total number of tissues. The entropy score for each peak across tissues can be defined as H = −1*sum(Ri * log2Ri) (1 < i < N), where the value of H ranges between 0 to log2(N). An entropy score close to zero indicates the accessibility of this peak is highly tissue-specific, while an entropy score close to log2(N) indicates that this peak is ubiquitously accessible^38^. Based on the distribution of entropy scores, peaks with score less than 3.5 were selected as tissue-restricted peaks.

We searched TF motifs in tissue-specific peaks using the *findMotifsGenome.pl* script of the HOMER^39^ version 4.9.1 software with default settings. We then generated a motif enrichment matrix^32^, where each row represents the P-value of a motif and each column represents a tissue. The 50 motifs with the top CV values and mean values greater than 20 were displayed.

### Code availability

The R codes used for correlation analysis, identification of tissue-specific chromatin accessible regions, and tissue-specific TFs are available in the supplementary materials (Supplementary File 1). A repository list containing the chromatin accessibility raw count matrix and the motif enrichment matrix is available online^32^.

## Data Records

A complete list of the 66 tissue samples is given in Table 1 and Table 2. All raw data have been submitted to the CNGB Nucleotide Sequence Archive^40^. The raw data have also been submitted to the NCBI Sequence Read Archive^41^. The ATAC-seq QC results and count matrixes have been submitted to the Figshare^32^.

## Technical Validation

### Data QC from the pipeline with IDR quality control

We evaluated our ATAC-seq dataset by a series of commonly used statistics, including the number of total read, mapping rate, the proportion of duplicate read, the number of mitochondrial read, and the number of final usable read (Table 2). For each replicate, we obtained an average of 78 million reads, which was previously shown to be enough for the detection of accessible regions^15,31^. In agreement with published ATAC-seq profiles^15^, the chromatin accessibility fragments show size periodicity corresponding to integer multiples of nucleosomes^32^ (Fig. 2b). The successful detection of accessible regions is also supported by the observation of strong enrichment of ATAC-seq reads around transcription start sites (TSSs)^32^ (Fig. 2a, d).

**Figure 2.**
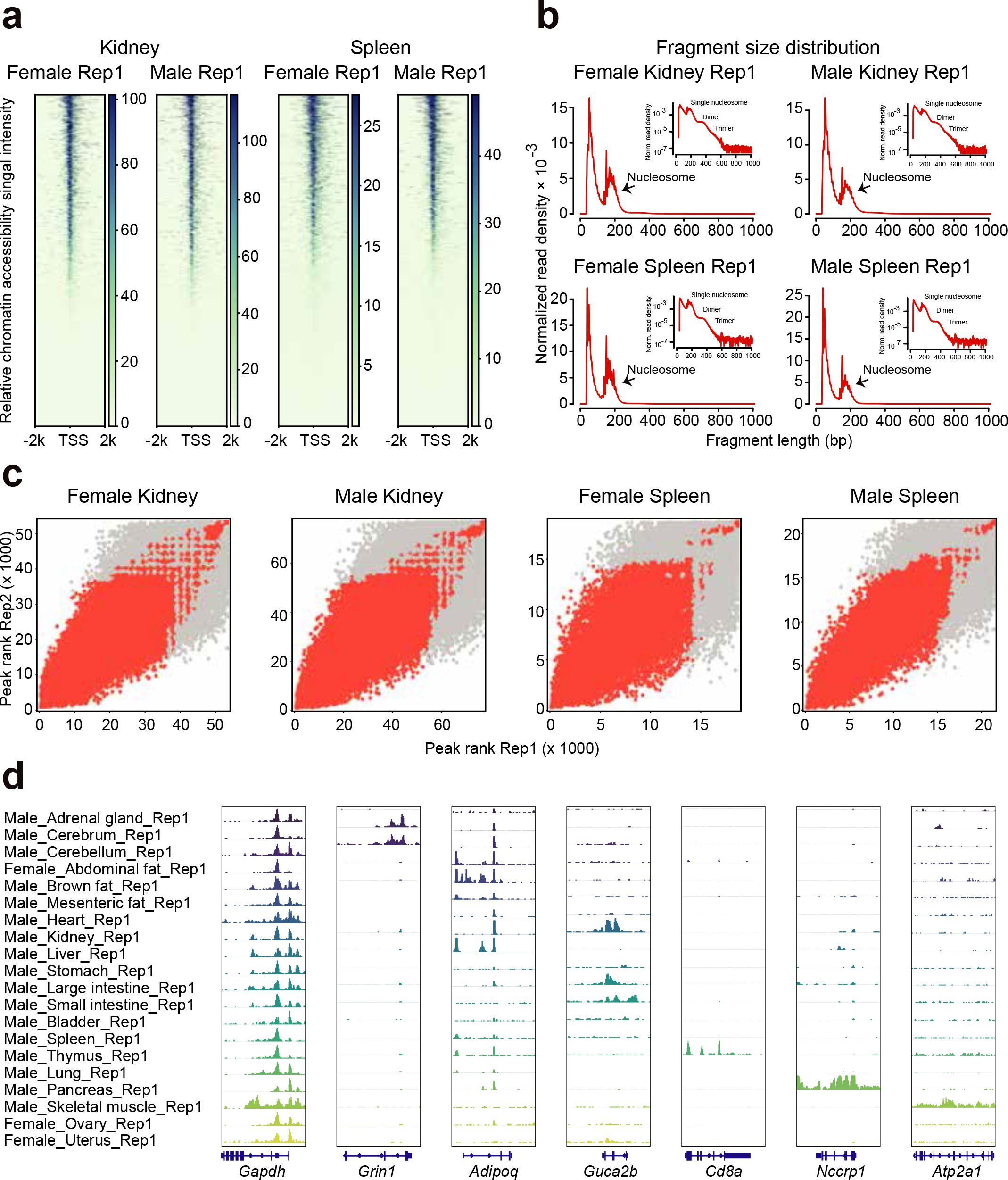
ATAC-seq data quality metrics. (**a**) The ATAC-seq signal enrichment around the tran-scription start sites (TSSs) for 4 representative samples (kidney or spleen of male or female mice) (**b**) The insert size distribution of ATAC-seq profiles for the same samples shown in 2a. (**c**) The irreproducible discovery rate (IDR) analyses of ATAC-seq peaks for the indicated samples. The scatter plots show points for every peak, with their location representing the rank in each replicate. (**d**) Genome browser views of ATAC-seq signal for the indicated housekeeping gene (Gapdh) and tissue-specific genes.

To evaluate the reproducibility of accessible element discovery between replicates, we identified accessible regions in both replicates by using the MACS2^33^ algorithm. We then applied the IDR method^34^ to find peaks that were reproducible between replicates in each tissue type (Fig. 2c). We identified an average of 43,421 reproducible peaks (Table 2). For downstream analyses, we filtered out low-quality data where the TSS enrichment scores are less than 10.0 X and the number of reproducible peaks are less than 10,000.

### Reproducibility of biological samples and comparison with published studies

The Pearson correlation analysis was used to further examine the reproducibility of biological and technical replicates. Heatmap clustering of Pearson correlation coefficients from the comparison of 66 datasets revealed a strong correlation between replicates of the same tissue (Fig. 3b), but a lower correlation between profiles from distinct tissues. This result is also supported by t-distributed stochastic neighbor embedding (t-SNE) analysis with tissue-restricted peaks of all profiles (Fig. 3a, c). Interestingly, correlations between replicates from mice of the same gender are generally higher than those from the opposite gender. This can be seen in the cerebrum where the correlation coefficient between replicates of female mice is 0.99 (Fig. 3d), while the coefficient between male and female is slightly reduced (0.96). We also compared our data to ATAC-seq profiles of postnatal mouse (day 0) tissues downloaded from ENCODE project^42,43^. Importantly, we found that both heart and lung were comparable with each other (Fig. 3e). Taken together, these analyses strongly suggest that our ATAC-seq profiles can reliably detect accessible chromatin regions in the mouse genome and can be used as a basic reference ATAC-seq dataset for future studies.

**Figure 3.**
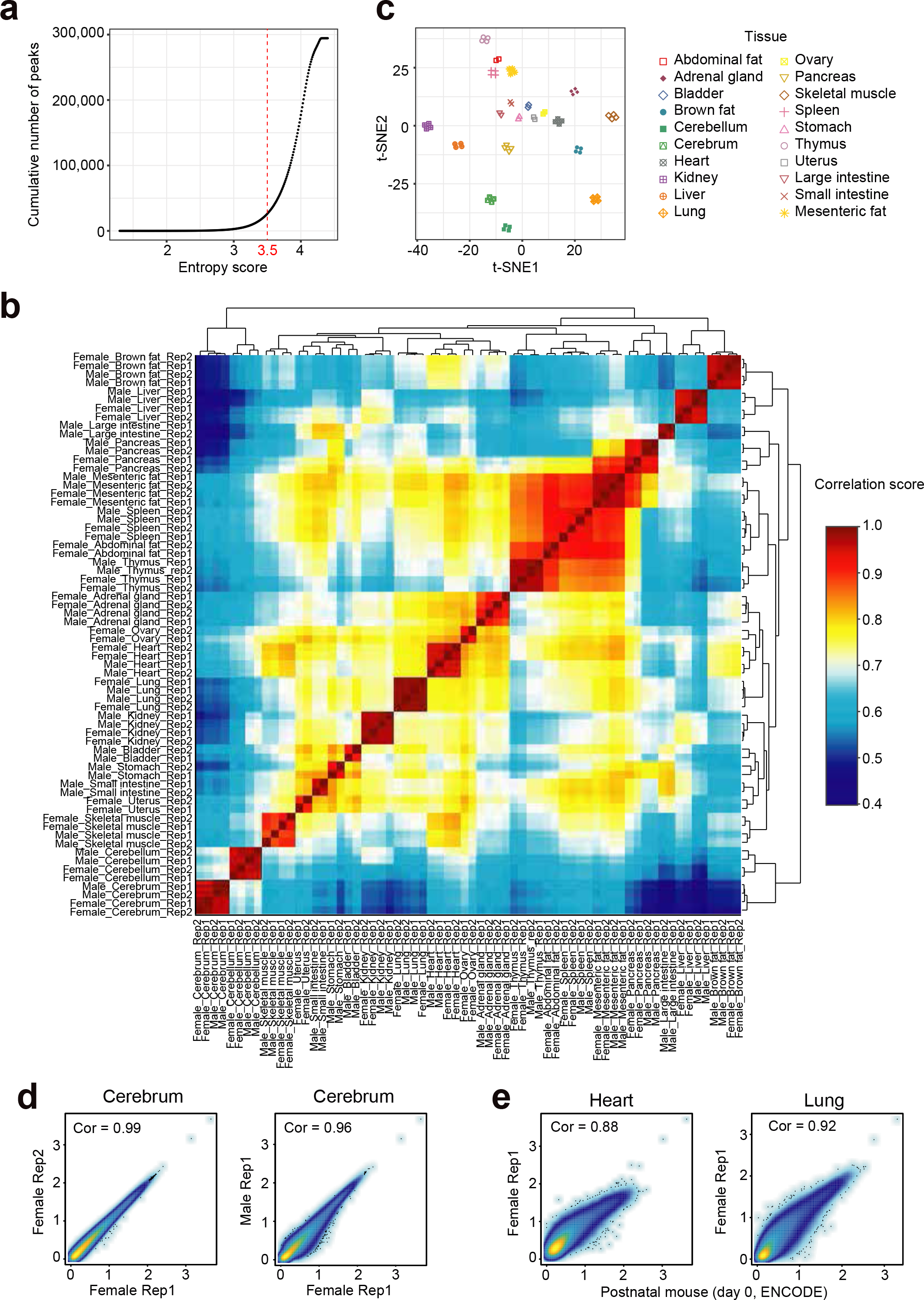
Evaluation of reproducibility across the ATAC-seq datasets. (**a**) The distribution curve of peak entropy scores. (**b**) Heatmap clustering of correlation coefficients across all 66 tissue ATAC-seq profiles. (**c**) t-SNE plot of all 66 ATAC-seq profiles based on the 26,916 tissue-specific peaks. (**d**) Scatter plots showing the Pearson correlations between technical (left) and biological (right) replicates. (**e**) Scatter plots showing the Pearson correlations between ENCODE postnatal mouse (day 0) datasets and our ATAC-seq profiles.

### Inferring tissue-specific transcription factors

In an effort to validate the tissue-specific TF motifs identified in our dataset we compared them to results from previous studies. Log2 RPM of the tissue-restricted peaks was shown in the heatmap (Fig. 4a). For example, we observed high enrichment of the NeuroG2 motif in cerebellum and cerebrum (Fig. 4b), in agreement with the role of NeuroG2 in controlling the temporal switch from neurogenesis to gliogenesis and regulating laminar fate transitions^23^. In brown fat, we found the CCAAT-enhancer-binding proteins (CEBP) motif to be highly specific (Fig. 4b). This is supported by a previous study demonstrating that CEBP can cooperate with PRDM16 to induce brown fat determination and differentiation^24^. In addition, other well-known tissues-specific motifs such as the liver-specific HNF family TF motifs (Hnf1, HNF1b, and HNF4a)^25^ and heart or skeletal muscle specific Mef2 family motifs (Mef2a, Mef2b, Mef2c, Mef2d)^26,27^ were validated in our study. To further validate whether the overall motif enrichment in each tissue can reflect tissue specificity we performed hierarchical clustering of tissues using Euclidean distance (Fig. 4c). This provided a result similar to hierarchical clustering of various mouse tissues based on RNA-seq data^44^. In addition, examination of tissues from the gastrointestinal (GI) tract (i.e., large intestine, small intestine, and stomach) showed tight clustering (Fig. 4c), which is likely due to their common functions such as lipid metabolism and energy hemostasis^45,46^. Skeletal muscle and heart tissue are found in the same branch, suggesting that patterns of chromatin accessibility in the two tissues are highly influenced by shared TF motifs such as those from the Mef2 family^45^.

**Figure 4.**
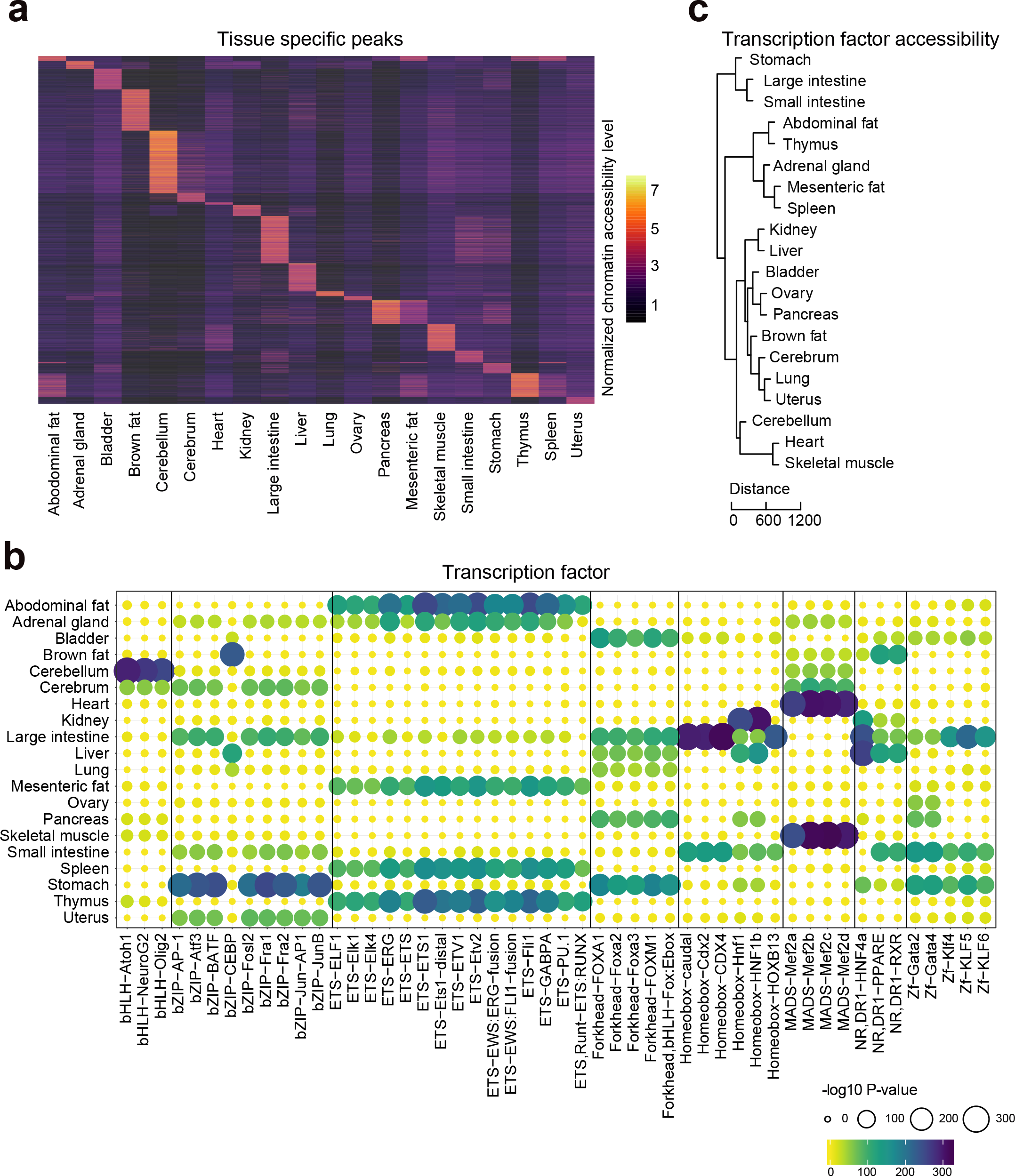
Identification of tissue-specific chromatin accessible elements and transcription factors. (**a**) Heatmap clustering showing the tissue-specific accessible elements. (**b**) Enrichment of the indicated TF motifs in each tissue. The size and color of each point represent the motif enrichment P-value (-log10 P-value). (**c**) The hierarchical clustering of transcription factor enrichment scores in each tissue. Euclidian distances are shown in the legend.

## Usage Notes

The ATAC-seq data processing pipeline, including read mapping, peak calling, IDR analysis, and read counting were run on the Linux operating system. The optimized parameters are provided in the main text. All R source codes used for the downstream data analyses and visualization are provided in Supplementary File 1.

## Acknowledgments

We thank all members of the Cell and Development Lab (BGI) for helpful comments and Jian Zhang and Jie Chen from Shenzhen Institutes of Advanced Technology for assistance with sample collection. This work was supported by the National Key R&D Program of China (No. 2016YFC1303902), the Shenzhen Municipal Government of China Peacock Plan (No. KQTD20150330171505310), and the Shenzhen Municipal Government of China (No. JCYJ20160531194327655).

## Author contributions

C.L., L.L., M.W. and X.W. conceived the idea. C.L. and M.W. collected samples. W.M. and C.L. generated the data. J.X., J.Xia, J.L., T.F., M.C. and Y.Y. assisted with the experiments. X.W. analyzed the data with the assistance of C.L., X.D., P.Z. and L.W‥ C.L. wrote the manuscript with the input of X.W. and M.W‥ L.L. and B.A.P. supervised the study and revised the manuscript. X.Z., F.C., W.Z, A.C. and Z.S. provided helpful comments on this study. All authors reviewed and approved the final manuscript.

